# “Devil’s stairs”, Poisson’s Statistics, and Patient Sorting via Variabilities for Oxygenation: All from Arterial Blood Gas Data

**DOI:** 10.1101/2021.08.10.455835

**Authors:** Gennady Chuiko, Yevhen Darnapuk, Olga Dvornik, Yaroslav Krainyk, Olga Yaremchuk, Anastasiia Diadenko

**Affiliations:** dept. of Computer Engineering, Petro Mohyla Black Sea National University, Mykolayiv, Ukraine

**Keywords:** Blood oxygen saturation, devil’s staircase, data mining, subsets, kernel density estimations, Poison distribution

## Abstract

This report deals with arterial oxygen saturation (SaO2) for healthy adults. A comparably small data set (20 persons) holds 3-minute records of SaO2. The sample rate was 200 Hz. The charts have the looks of a “devil’s stairs.” A few (from 1 to 10) detectable oxygenation levels form the stair’s treads, more or less long. “The risers” have two types (up and down), and all have virtually the same height, about 1 %. The inter-level shifts ( 0 to 42 switches per record) turned out a rare event at the actual sample rate. The number of switchings meets the Poisson distribution. There were found three visibly varied intensities for the switch-overs within the data set. Histograms also show the co-existing of no fewer than three subsets into the data set. The subsets differ by the intensity of switch-overs, amounts of possible levels, relative frequencies of most probable levels (modes), etcetera. In short, those all are diverse variability quantifiers. The higher variability subset has about 25 %, the lower one - 45%.

## I. Introduction

Arterial Blood Gas (ABG or SaO2) analysis is one of two used in clinical practice methods of blood oxygen saturation measuring [1]. It is direct but invasive, more laboring, and pricey than another one – peripheral Pulse Oximetry (SpO2). Still, many authors prefer the SaO2 as a more reliable way of the blood oxygen lack (hypoxia) finding [2]

Blood oxygenation got a particular weight during the coronavirus (COVID-19) pandemia. This disease’s early stages are tied with “silent hypoxia” due to poor saturation [3, 4]. It means the oxygen lack (hypoxia) might be almost stealth for the level around 90 % and below. The norm is in the range of 95-100% for healthy adults [4].

Reference [5] recently reported about three (at least) subgroups that co-exist among healthy adults. They differ by oxygen saturation variabilities. The authors have used the SpO2 data set and Recurrence analysis there. Presumed that the people with higher variability might be a riskier fraction to COVID-19.

Here we intend to test the SaO2 data set. The methods of analysis also will be rotated. Thus, we aimed to check the results of [5] and confirm them if it turns out to be possible.

## II. Data and methods

### A. Data origins and main features

We consider SaO2 data for the small set (n=20 anonymous subjects with IDs from 1 to 20) randomly picked up from the vaster database [6, 7]. The database was contributed in 2018 by two Massachusetts General Hospital’s (MGH) laboratories: Computational Clinical Neurophysiology Laboratory and the Clinical Data Animation Laboratory.

The database includes N=1,985 subjects who were monitored at MGH. The data is partitioned into a balanced training set (N1 = 994) and test one (N2 = 989). The reference [8] holds a detailed report about this database.

We have been focused on relative short-time (3-minute length) SaO2 records. The sampling rate was equal to 200 Hz [6, 7]. Thus the 3-minute size of a signal means 36,000 counts per record. A pretty long one from this point of view is not?

Unfortunately, the resolution, or at least the device’s accuracy, is pointed out nowhere in these sources. At all, the accuracy of oximeters is a feature that is rarely indicated [9]. We will return to this point later, and the oximeter’s resolution will be estimated, but implicitly.

### B. Methods

This study’s main methods are statistical methods collected in the extensive software package “Statistics” of Maple 2020 [10]. In rare cases, we have written short programs (procedures) in Maple language. The plotting of the Recurrence plots had needed it. Another case was the counting of the number of singularities in a record.

Besides, we relied on the kernel density estimations (or the KDE method [11,12] ). One can consider KDE as the way of histogram smoothing also. Some simple statements of the theories for singular multifractal functions [13] and the “devil’s-staircases”-like behavior for time series [14] were in use too. The Recurrence plots [5] only illustrate and confirm the results.

Subgroups with low, middle, and high variability of saturation were marked in [5] as “strong,” “middle,” and “weak.” We will keep on this labeling below.

## III. Results

### A. “Devil’s staircases” and switch-overs between nearest possible saturation levels

We will show only three of the twenty subjects of the data set to save space. These three belong to different) subgroups, as we would like to argue below.

The step-like shape of signals is typal for the singular functions, which have zero-equal derivatives almost everywhere. “Almost” means the exception for a set of points, where the functions can have the discontinuities of the first kind, also called jump- or step-like ones. Those sets of points have the zero-measure and are the fractals by themselves [13]. The reader can see these point sets on the lower row of Fig. 1. Math functions, having described above singularities, are termed as “devil’s staircases.” [13, 14],

**Fig.1.**
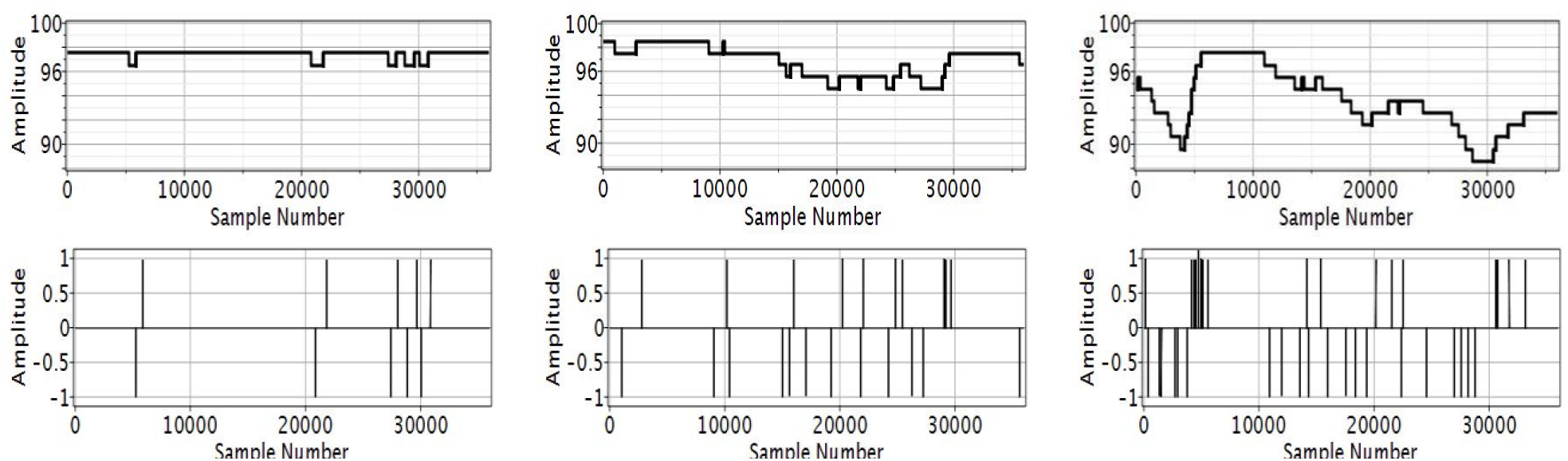
Arterial blood oxygenation (SaO2) records for subjects with following IDS: {1, 13, 3} from left to right (the upper row). The difference between sequential terms of upper series: SaO2(n+1) - SaO2(n) where n is sample number (the lower row). Positive and negative differences segments indicate the up and down steps of upper graphs. Their hights show the hights of steps (switch-overs amolitudes). Amplitudes are shown in percents.

The number of step-like breaking varies from 0 to 42 for the different subjects and is an integer. For instance, it was 10, 22, and 37 for the examples of Fig.3. This number can be a measure of variability by itself. As more significant is this number, the more large variability is.

**Fig.2.**
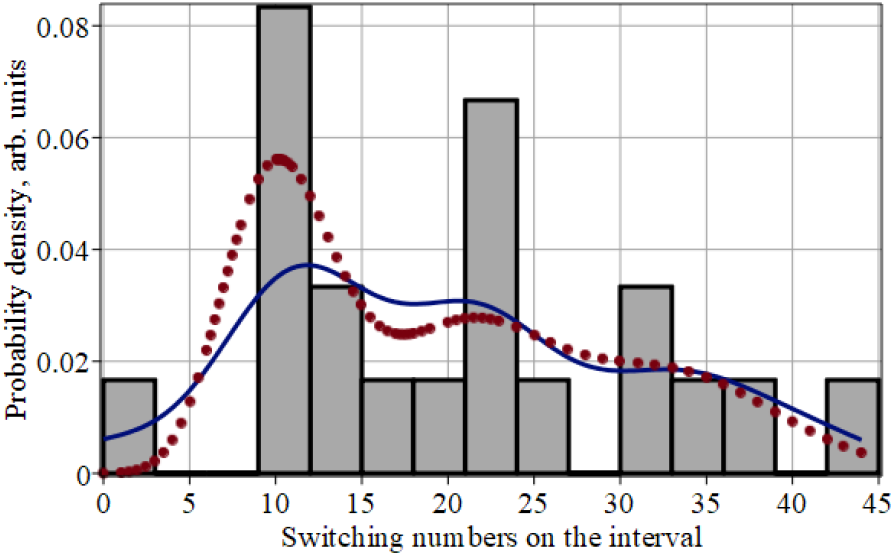
Histogram (bars) for the switching amounts on the 3-minute interval, KDE probability density estimations (solid line), and three-modal Poisson distribution’s weighted model (dot line). Weights’ vector is {0.45, 0.30, 0.25}

**Fig.3.**
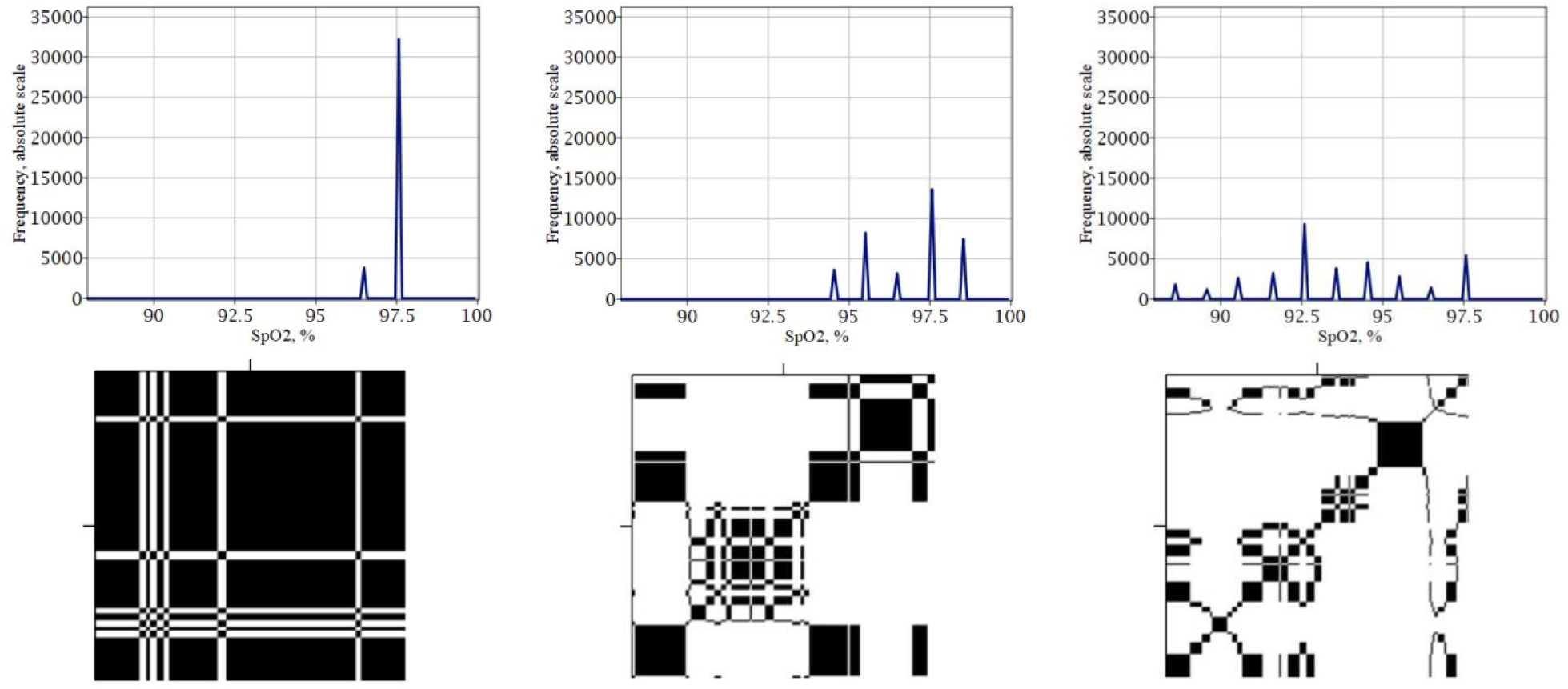
Spectra of absolute frequencies of possible saturation levels (upper row) and Recurrence Plots (lower row) for subjects with IDs: {1,13, 3}. The Recurrence Ratios are: {0.812, 0.256, 0.140} respectively. Note the visual differences among left, middle and right graphs.

Note, the heights of steps are almost the same for the various records and are unchanged—the only exception was for the subject with ID=13. There detected sole double-switching with a height of about 2 %. The resolution of the measuring device causes such an identity of steps (switch-overs). The current sensitivity threshold was most likely touch lower than the hights of steps. Hence, the closer levels, if they existed, just were unresolved.

The step-like switchings are discrete and relatively rare occasions, comparing their quantity with the number (N=36000) of trials. Noteworthy is following:

- Each switching occurs independently of other ones.
- The average rate at which switching occur is independent of any other switchings.
- Two switchings cannot occur at precisely the same instant.

The above conditions define the number of switching on the recording interval as Poisson distributed discrete random value. Let us verify this by constructing the histogram of these numbers what is showed in Fig.2

The histogram is three-modal, as well as its KDE smoothing curve (solid line). Hence, the co-existing of three subgroups inside our data set manifests itself. Subgroups differ by the average rate of switchings occurrence. If we take the 3-minute interval, then these average rates are the following: 11 ± 3, 22 ± 5, and 34 ± 6 respectively for strong, middle, and weak subgroups. The fractions (weights) of subgroups are 0.45, 0.30, and 0.25 in the same order.

These estimations allow the build weighted Poisson distribution for the entire data set. This simple model, showed by the dotted line in Fig. 2, turned out close enough to KDE estimations (solid line). So, the stratification of our data got the proofs.

### B. Frequencies and Recurrence plots

Consider the upper left graph of Fig.1. There all is simple and clear. Only two possible saturation levels exist on the interval. The higher of them cover most of the interval, while the lower one - much lower part. Let call the cumulative amount of trials that manifest itself that or another possible level as their absolute frequencies. If these frequencies are normed on the total number of tests (N=36000), we deal with relative frequencies. The higher are these last, as higher is the probability of the saturation level and its recurrence ability. The level with the highest frequency is the mode.

Fig.3 shows the absolute frequencies plots and Recurrence Plots [15] for the same subjects and in the same order as in Fig.1. The strong subset subjects (7 persons) had 1 to 3 possible saturation levels with clearly dominated powerful modes (left upper graph of Fig.1). The weak subset (5 persons) displays from 6 to 10 possible levels with many low-rise modes. Besides, the Recurrence Ratios are much higher for the subjects of the strong subset. That show the lower row of Fig.3

### C. Most probable oxygen saturation levels (modes) and their relative frequencies

Fig. 1 pointed out a positive correlation between mode frequencies and repeatability coefficients, which have been successfully used to measure variability [5]. Indeed we found the correlation coefficient between Recurrence Ratios and relative frequencies of modes as positive and statistically significant: 0.985. Hence the relative frequencies of modes can be the once more measure of the variability. Consider this new measure in Fig.4, where the histogram and its KDE smoothing line are presented. Both are multimodal.

The medians for subsets are 76, 40, and 31 percent from strong subset to weak one. Thus, this descriptor good for the strong subset resolving, but a few worse works between the other two. The descriptor from the above section seems to be preferable in this sense.

**Fig,4.**
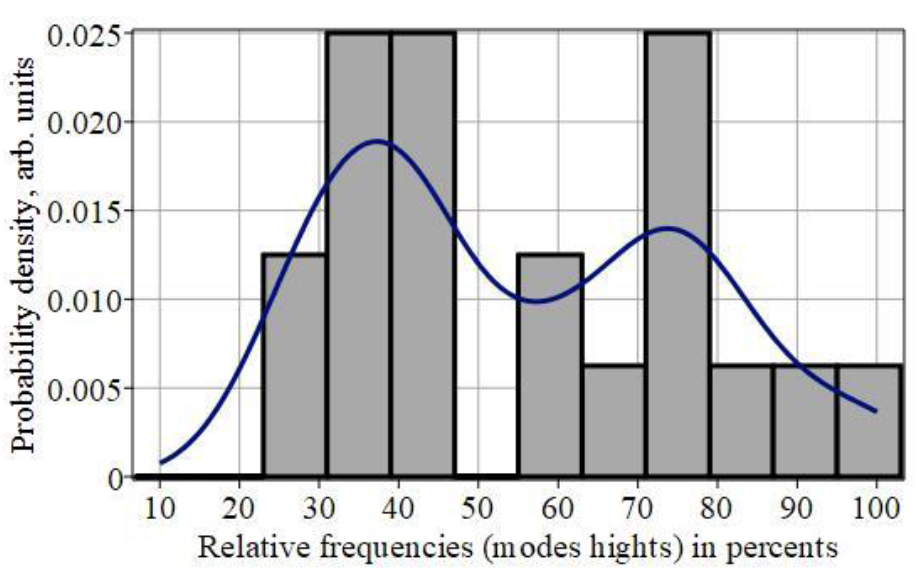
Histogram (bars) and KDE smoothing line for the modes relative frequencies.

### D. Comparison between two suggested variability

#### descriptors

Compare their box-whiskers-plots in Fig.5. The heights of boxes show the interquartile ranges, the whiskers - the entire ranges, the horizontal segments inside boxes are medians, and points show outliers if they are. Pay attention to the particular overlapping of two subsets ranges on the left-hand side graph in Fig.5. The right-hand chart has no overlappings.

**Fig,5.**
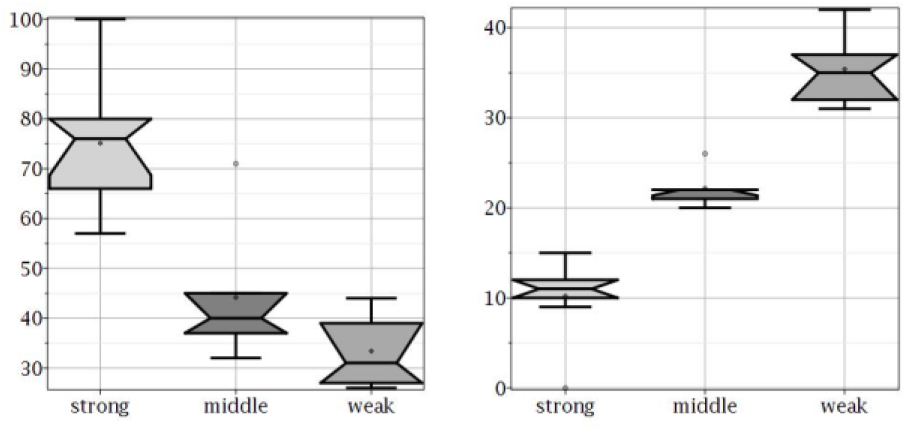
Statistical box-plots for relative frequencies of modes (left habd side) and for numbers of switch-overs on 3-minute interval (right hand side ) builded for three subsets.

Finally, let consider the so-called probability-probability plot for the switchings numbers. Such graphs show the deviations of a factual distribution from the Gauss normal one. Fig. 6 shows the distribution for the switch-overs numbers is close enough to Gauss one line. Notably, the points form a few steps with equal heights. One can see a typical “devil’s staircase” again, but now it is mono-fractal

**Fig. 6.**
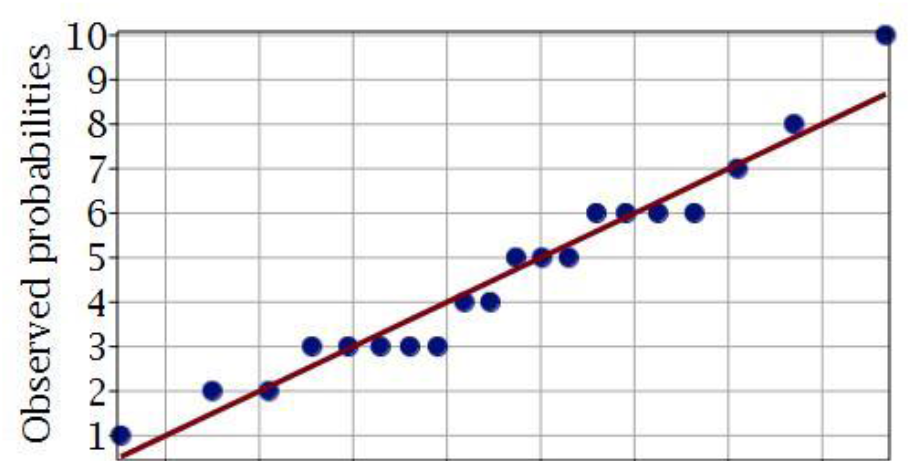
The normal plot for the switch-overs numbers. The thin line presents th Gauss distribution, the points a factual one.

## IV. Discussion

This report claims about three subsets, differing by the variability of SaO2. That meets three subsets of [5] where SpO2 data has been studied. The sums of weights are equal to one, to be sure. One more common trait is the weak subset’s lowest weights concerning two others, but no more.

Still, the weights of the same subsets are various (see Fig.7). We cannot explain now the reasons for divergences unambiguously. Most likely, for the exacter estimations of the subsets probabilities, we need extended data sets, while 36 in or 20 subjects in our study are maybe not enough for this purpose. Perhaps also, there are some differences in these weights for SaO2 and SpO2 data. The various methods of variability evaluations in [5] and here also can have significance.

**Fig.7.**
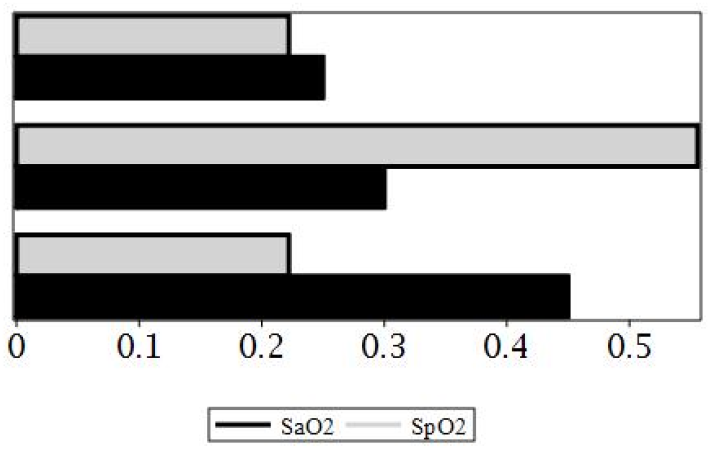
Bar chart for the weights (probabilities) of thee subsets inside data sets. The weights vector components shown as black for SaO2 data and as gray for SpO2 data [5] pairwise.

## V. Conclusions

The results of the study allow a few conclusions:

- A few almost equidistant discrete oxygen saturation levels, which occupy the relatively narrow range, are observable for each subject. There might be from 1 to 10 levels for a 3-minute interval.
- The levels have various probabilities. The maximal ones might be from 0.31 up to 1.
- The switch-overs between possible levels are the rare (low-frequencies) events, which meet the discrete Poisson’s distribution. The average frequencies for the three subsets are: {0.057, 0.123, 0.197} Hz.
- The difference of mean switching frequencies among subsets is statistically significant (p<0.0012 at confidence 0.99). We confirm the results [5] got for SpO2 data afore, using independent methods at that.
- The switching frequencies suggested here descriptors of the variability need much less computing labor than one offered in [5.]

## Acknowledgment

This report is a part of the research project “Development of hardware and software complex for non-invasive monitoring of blood pressure and heart rate of dual-purpose” with registration number 0120U101266. Ukrainian Ministry of education and science financially supports this project, and the authors are grateful for that.

